## Supplemental Data 1 for "Afadin-deficient mouse retinas exhibit severe neuronal lamination defects but preserve visual functions"

| **Primer name** | **Sequence (5' to 3')** |
| --- | --- |
| Opnsw Fw | CAGCATCCGCTTCAACTCCAA |
| Opnsw Rv | GCAGATGAGGGAAAGAGGAATGA |
| Opnmw Fw | CTCTGCTACCTCCAAGTGTGG |
| Opnmw Rv | AAGTATAGGGTCCCCAGCAGA |
| Nrl Fw | GCTGTGCCTTTCTGGTTCTGA |
| Nrl Rv | GCTCCCGCTTTATTTCGAACT |
| Rho Fw | GACTCTGCCAGCTTTCTTTGCT |
| Rho Rv | GCGTCGTCATCTCCCAGTGGA |
| Trpm1 Fw | ATGCGCCCATTGTCAAGTTC |
| Trpm1 Rv | TTCTCCAATGCAAGGCTCACA |
| Grm6 Fw | GTCCATCATGGTCGCCAATGT |
| Grm6 Rv | AGTCATAGCGTGTGGAGTCAC |
| Chx10 Fw | GGCGACACAGGACAATCTTTA |
| Chx10 Rv | TTCCGGCAGCTCCGTTTTC |

**Supplementary File 1 The primer sequences used for RT-qPCR**
